# BLEND: Probabilistic Cellular Deconvolution with Automated Reference Selection

**DOI:** 10.1101/2024.08.02.606458

**Authors:** Penghui Huang, Manqi Cai, Chris McKennan, Jiebiao Wang

**Affiliations:** Department of Biostatistics, University of Pittsburgh, De Soto St, Pittsburgh, 15261, PA, USA; Department of Statistics, University of Pittsburgh, S Bouquet St, Pittsburgh, 15213, PA, USA

**Keywords:** Cellular deconvolution, Single-cell RNA sequencing, Bayesian estimation, Gibbs sampling, EM algorithm, Maximum a posteriori estimation

## Abstract

Cellular deconvolution aims to estimate cell type fractions from bulk transcriptomic and other omics data. Most existing deconvolution methods fail to account for the heterogeneity in cell type-specific (CTS) expression across bulk samples, ignore discrepancies between CTS expression in bulk and cell type reference data, and provide no guidance on cell type reference selection or integration. To address these issues, we introduce BLEND, a hierarchical Bayesian method that leverages multiple reference datasets. BLEND learns the most suitable references for each bulk sample by exploring the convex hulls of references and employs a “bag-of-words” representation for bulk count data for deconvolution. To speed up the computation, we provide an efficient EM algorithm for parameter estimation. Notably, BLEND requires no data transformation, normalization, cell type marker gene selection, or reference quality evaluation. Benchmarking studies on both simulated and real human brain data highlight BLEND’s superior performance in various scenarios. The analysis of Alzheimer’s disease data illustrates BLEND’s application in real data and reference resource integration.

## 1 Introduction

Bulk RNA sequencing (RNA-seq) technology provides a valuable approach to studying gene expression patterns across different tissues or conditions [1]. However, bulk RNA-seq is typically performed on tissues composed of many cell types, where estimating the proportion of each cell type in each sample can enrich analyses and help address biases that occur due to cell type heterogeneity [2, 3].

Several biochemical pipelines have been developed to estimate cell type proportions and account for the aforementioned biases, including directly measuring cell type fractions or cell type-specific (CTS) expression (e.g., single-cell RNA-seq). Unfortunately, the high cost of these pipelines precludes their use in large-scale studies. Consequently, many *in silico* cellular deconvolution methods have been developed in recent years to estimate cellular fractions as a computational alternative. Cellular fraction estimates allow analysts to infer how cell type proportions vary with different phenotypes, account for cell type confounding, and facilitate downstream CTS analyses, such as estimating CTS gene expression, CTS differential expression, and CTS expression quantitative trait loci analyses [4, 5].

Cellular deconvolution models bulk gene expression as the weighted average of CTS gene expression, where the weights are cell type proportions. Among existing methods to estimate cell proportions, reference-based methods have shown more robust performance compared to unsupervised reference-free methods [6, 7]. Reference-based methods rely on single or sorted cell data to construct a CTS expression reference matrix whose entries give each gene’s expected expression in each cell type. Due to the challenges of incorporating inter-subject variation in CTS expression into deconvolution models, existing methods assume the same reference matrix shared across all study subjects and provide no guidance to choose appropriate references. This belies the fact that there is often substantial heterogeneity in CTS expression that arises due to differences in experimental conditions, technical batch effects, and variation in reference sequencing technologies. Nonetheless, the choice of reference is the most important factor affecting deconvolution accuracy [7–9]. It is therefore critical to develop new statistical methods to personalize signature matrices for each subject to account for inter-subject variation in CTS expression.

To address the above issues, we developed BLEND, a hierarchical Bayesian model that is, to our knowledge, the first deconvolution method to leverage multiple available references to create personalized references for study subjects. Inspired by Latent Dirichlet Allocation (LDA) [10], we use a “bag-of-words” representation for bulk RNA-seq count data whereby we view a read count as a word, a cell type as a topic, and a bulk sample as a document. In this analogy, a cell-type topic’s reference is its distribution over words. However, unlike conventional LDA-based deconvolution methods, which assume references are shared across bulk samples, BLEND allows references to be sample-specific and uses the data to learn each sample’s most appropriate reference among all possible references in the convex hull of available references. We show through extensive realistic simulations and real data applications that such reference learning drastically improves cell type proportion estimates. BLEND also stands out for being user-friendly, as it eliminates the need for manual reference evaluation, data transformation and normalization, and the selection of cell type marker genes.

## 2 Results

### 2.1 Overview of BLEND method

BLEND is a cellular deconvolution method that can leverage multiple references as visualized in Fig. 1. First, BLEND individualizes the most suitable reference matrix for each bulk sample by exploring convex hulls of available CTS reference vectors. Second, BLEND uses individualized references to deconvolve bulk count data by employing a “bag-of-words” representation. BLEND adopts a unified hierarchical Bayesian framework to incorporate these two steps. Two consistent parameter estimation strategies are provided: Gibbs sampling and EM-MAP. The detailed statistical model of BLEND is provided in Methods, and the algorithm derivation is in Supplementary Notes.

**Fig. 1.**
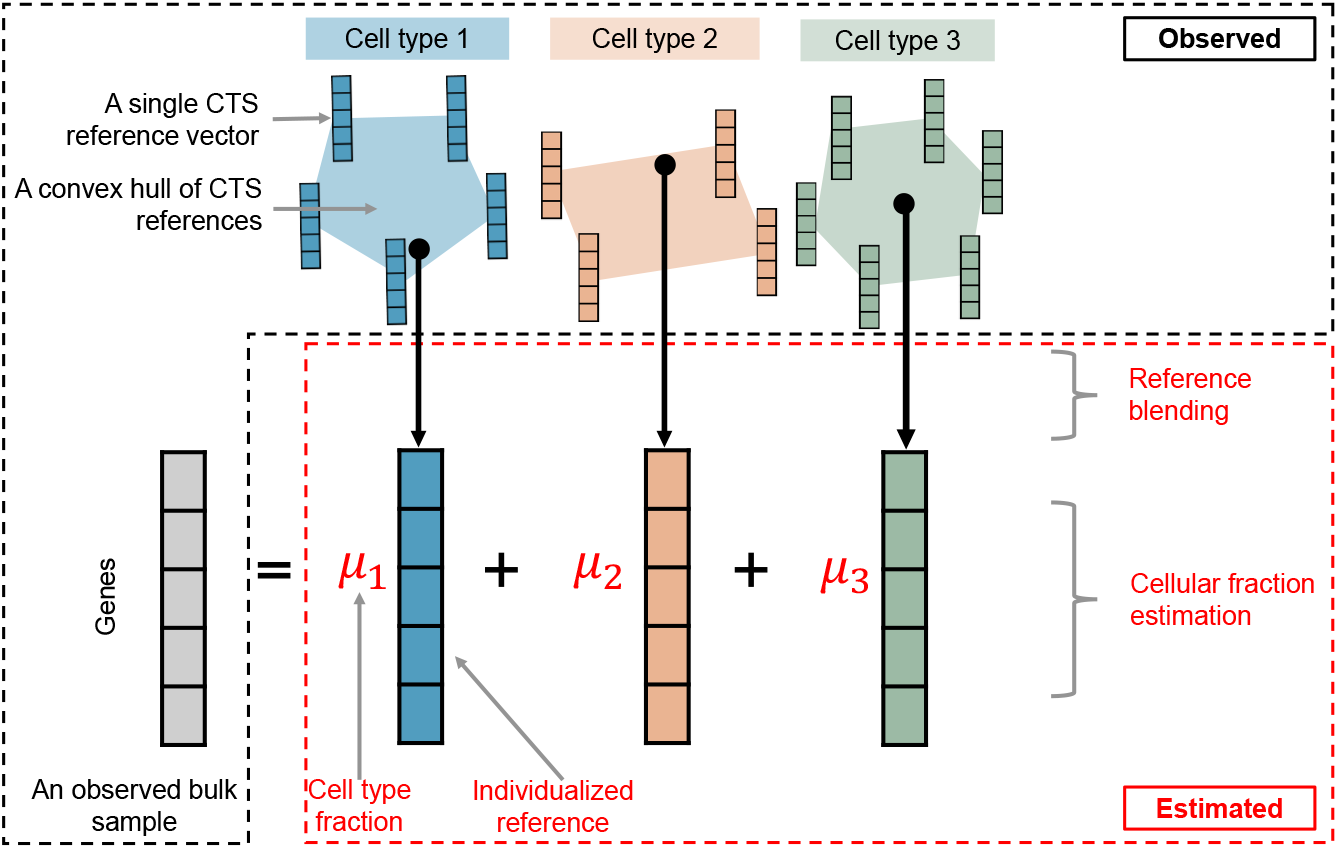
General framework of BLEND. BLEND takes multiple CTS expression references as input to deconvolve the targeted bulk count RNA-seq data. BLEND models each bulk sample as the weighted average of CTS gene expression, where the weights are cell type proportions (***µ***). For each bulk sample, BLEND explores its most suitable references in the convex hulls of available CTS references and performs deconvolution using a “bag-of-words” representation.

### 2.2 BLEND improves performance with partially matched references

In the BLEND model, we assume the CTS expression of bulk samples lies in the convex hulls of the provided references. We first tested BLEND with an oracle simulation following the generative process and confirmed BLEND’s accuracy in reference selection and cellular fraction estimation (Supplementary Notes). However, in most real application settings, this usually is not the case. Here, we used a real dataset to simulate data where only partial references were available.

Mathys et al. [11] collected 427 individuals’ postmortem dorsolateral prefrontal cortex (DLPFC) tissues from Religious Orders Study and Rush Memory and Aging Project (ROSMAP) [12]. They measured gene expression levels across 2.3 million nuclei isolated from these tissues using droplet-based single nucleus RNA-seq (snRNA-seq). Among all donors, 418 of them contained cells of six major cell types in the brain: astrocyte (astro), microglia (immune), inhibitory neuron (inh), oligodendrocyte progenitor cell (OPC), oligodendrocyte (oligo) and excitatory neuron (ex). To generate more realistic data in this simulation experiment, we randomly chose 40 out of 418 individuals (around 10%) to serve as references to deconvolve all 418 individuals. We generated pseudo-bulk samples by simply summing up counts of snRNA-seq data for each individual and used cell counts of different cell types to calculate the ground truth for cellular fractions of all simulated bulk samples.

We compared BLEND with four deconvolution methods: BayesPrism [13], which is a bag-of-words-based Bayesian method using a fixed reference; MuSiC [14], a weighted nonnegative least squares method that utilizes multi-subject references; Bisque [15] and hspe [16], both of which were recommended for brain tissue deconvolution by a recent benchmarking paper [17].

Pseudo-bulk samples generated from the 40 individuals were provided with the matched reference list, from which BLEND selects references. For each cell type, we calculated Lin’s concordance correlation coefficient (CCC) [18] between the true and estimated fractions across individuals. CCC ranges between minus one and one and measures the deviation from the line *y* = *x*, where values close to one indicate points lie closer to the line. CCC is preferred over other metrics like Pearson’s correlation because it combines the mean difference and correlation between two sets of points to capture both location and scale shifts from the ground truth. We also averaged CCC across cell types to get mean CCC (mCCC). As visualized in bar charts in Fig. 2a, BLEND has the largest CCC for each cell type. Moreover, BLEND selected matched references for all cell types (Fig. 2b).

**Fig. 2.**
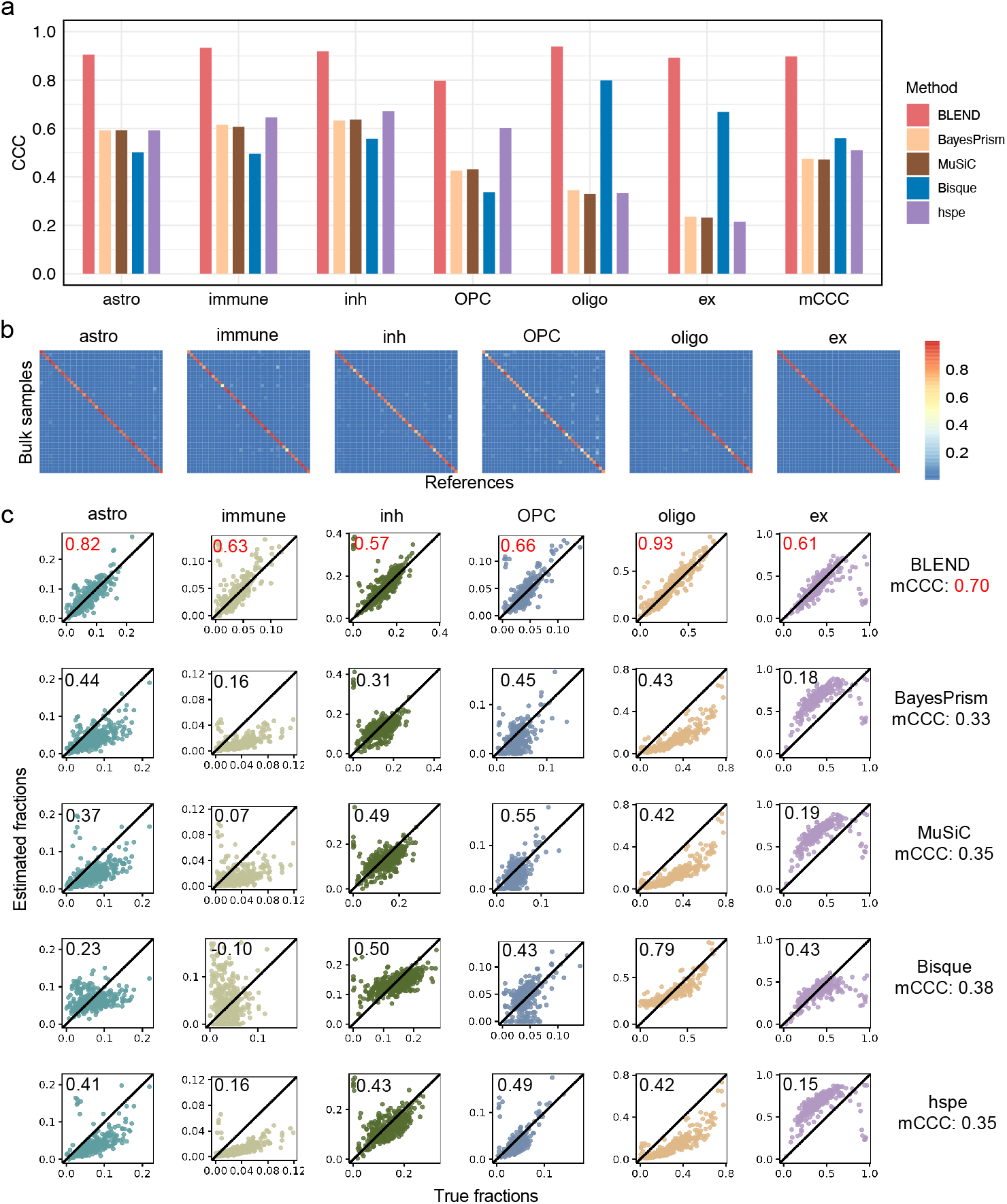
Benchmarking cellular deconvolution methods with partial reference simulation. **a**, Barchart visualization of CCC between the estimated and ground truth fractions for each cell type across 40 pseudo-bulk samples with matched references and mCCC across cell types. **b**, Heatmaps of estimated reference mixing proportions of the pseudo-bulk samples with matched references. Rows represent bulk samples, and columns represent references. **c**, Scatter plots of 378 samples with no matched reference. The x-axis represents true cellular fractions, and the y-axis corresponds to the estimated ones. CCC calculated for each cell type between estimated and true fractions is presented on the top-left corner of each plot, and mCCC across cell types of each method is listed on the right. The CCC of the best performer for each cell type is highlighted in red. astro: astrocyte; inh: inhibitory neuron; OPC: oligodendrocyte progenitor cell; oligo: oligodendrocyte; ex: excitatory neuron.

The more challenging case is the other 378 pseudo-bulk samples that do not have matched references. Natural discrepancies between pseudo-bulk data and references exist due to batch effects and biological differences among individuals. Fig. 2c plots each method’s estimated fractions against the simulated ground truth proportions. BLEND clearly has the largest CCC for each cell type, indicating BLEND’s personalized references substantially improve cell type proportion estimates.

### 2.3 BLEND alleviates the discrepancy between bulk and reference data

In most cases, the bulk data to be deconvolved do not have matched single-cell data. Thus, we usually use single-cell data from other datasets. However, the cross-data discrepancy in CTS gene expression is non-trivial. Here we show an example of two large-scale snRNA-seq studies collected from the same subjects. Similar to the Mathys data [11] we used in the simulations, Fujita et al. [19] collected snRNA-seq samples from the same study (ROSMAP), while the CTS expression of two datasets shows significant batch effects that form two distinct groups in the visualization (Fig. 3a). There are 231 individuals who have snRNA-seq measurements from both datasets. We designed two sets of cross-data simulation studies (Fig. 3b). The first set used Mathys scRNA-seq data as references to deconvolve pseudo-bulk data generated by collapsing Fujita snRNA-seq data, which is named Mathys-Fujita. Vice versa, the second set employed Fujita data as references to deconvolve pseudo-bulk data generated using Mathys data, which is named Fujita-Mathys. The simulation method was consistent with that in section 2.2.

**Fig. 3.**
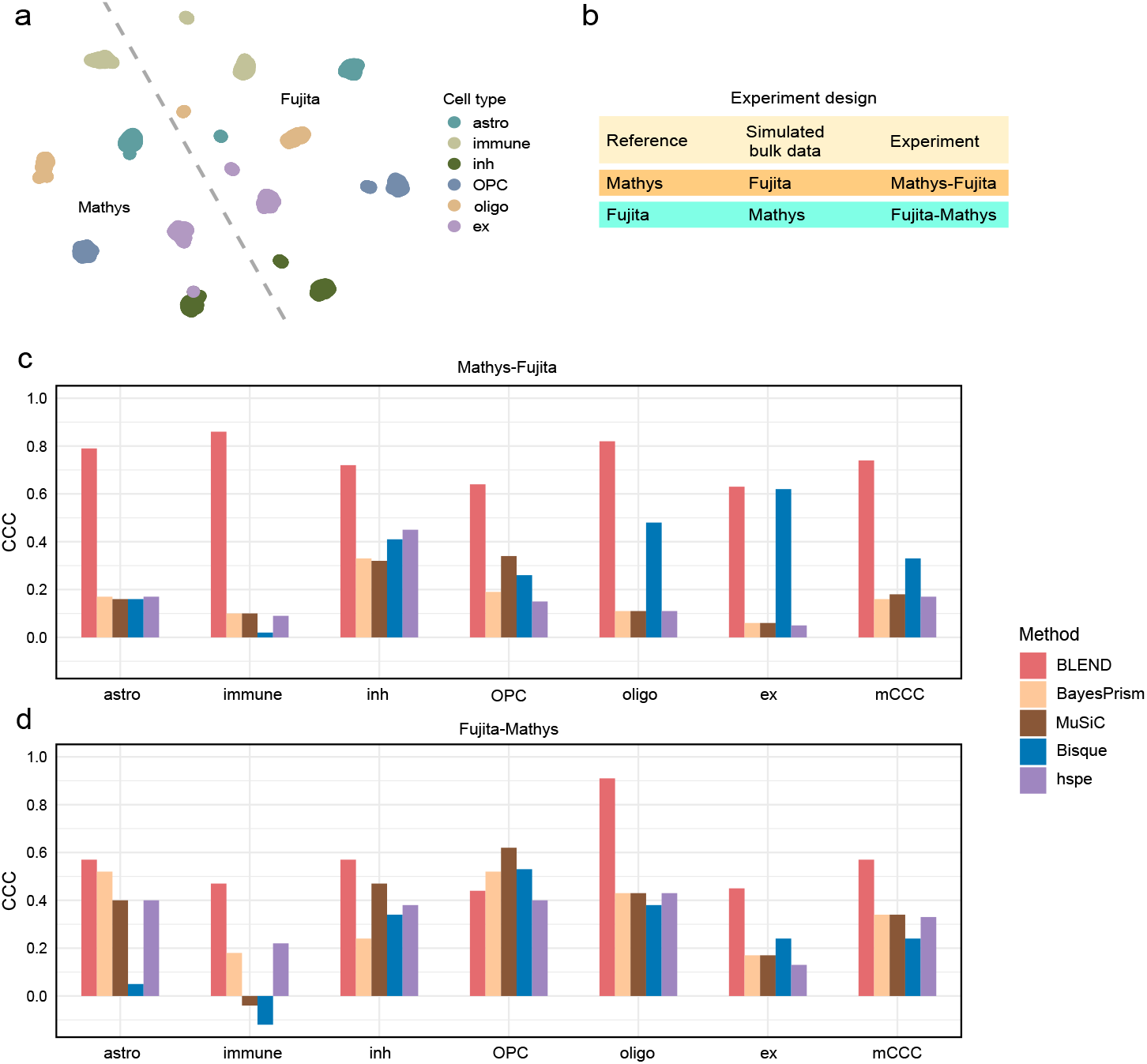
Benchmarking cellular deconvolution methods with cross-data simulation. **a**, UMAP of individual CTS expression of Mathys and Fujita shared individuals. **b**, Experiment design of cross-data simulation. **c-d**, Barchart visualization of CCC between the estimated and ground truth fractions for each cell type across all bulk samples and mCCC across cell types in the cross-data simulation. astro: astrocyte; inh: inhibitory neuron; OPC: oligodendrocyte progenitor cell; oligo: oligo-dendrocyte; ex: excitatory neuron.

After deconvolution. we calculated CCC between the estimated and ground truth fractions for each cell type across all bulk samples and mCCC in each simulation setting (Figs. 3c, and d). In the first setting, BLEND showed superior performance in all cell types, with an mCCC of 0.41 higher than the runner-up (Bisque). In the second setting, BLEND performed the best in five cell types and had the highest mean CCC.

### 2.4 BLEND provides accurate estimates in real data application

In the human brain, most studies use frozen postmortem tissues for RNA sequencing. Because cells are lysed, only nuclei can be measured using snRNA-seq techniques, while cytoplasm transcripts are usually left out [20]. Instead, bulk RNA-seq extracted total RNA from both nuclei and cytoplasm. Moreover, in real applications, reference data are often from individuals with varying phenotypes, different brain regions, or even different species than the bulk data [21]. Given these reasons, it may be unrealistic to assume there is no discrepancy between the CTS gene expression underlying all bulk samples and a specific reference.

To benchmark deconvolution methods in human brain, Huuki-Myers et al. [17] provided a multi-assay dataset of postmortem human DLPFC, including snRNA-seq, bulk RNA-seq, cell type fractions measured by RNAScope/Immunofluorescence (RNAScope/IF). They collected 19 tissue blocks from the anterior, middle, and posterior parts of DLPFC from 10 neurotypical adults. Different multi-assay technologies were performed on adjacent sections of the same tissue block. Before performing bulk RNA-seq, they used three RNA extraction methods: nuclear (Nuc), cytoplasmic (Cyto), and total (Bulk), combined with two library preparation techniques: polyA and RiboZeroGold. This resulted in a total of six library types. Finally, 110 bulk samples passed the quality control and were provided in the dataset. For RNAScope/IF, cellular fractions of six cell types were measured, including astrocyte, microglia, oligodendrocyte, endothelial, inhibitory neuron, and excitatory neuron. We then used the individual CTS expression from snRNA-seq as references to deconvolve bulk data. For the cellular fractions measured by RNAScope/IF, we found out that the proportions of cell types were systematically under-measured since the average total proportion was only 0.74, far less than 1. Thus, CCC or mean absolute error is not a proper metric here. Instead, we used Pearson’s correlation coefficient to measure the correlation between cellular fraction estimates and RNAScope/IF measurements.

We calculated the correlation between *in silico* estimates and RNAScope/IF measurements across cell types and bulk samples under different library types and RNA extraction combinations (Fig. 4). BLEND demonstrated unparalleled robustness across all data preparation conditions, outperforming all competing methods in five distinct settings, except in one setting it achieved the second-highest performance that was very close to the best performance.

**Fig. 4.**
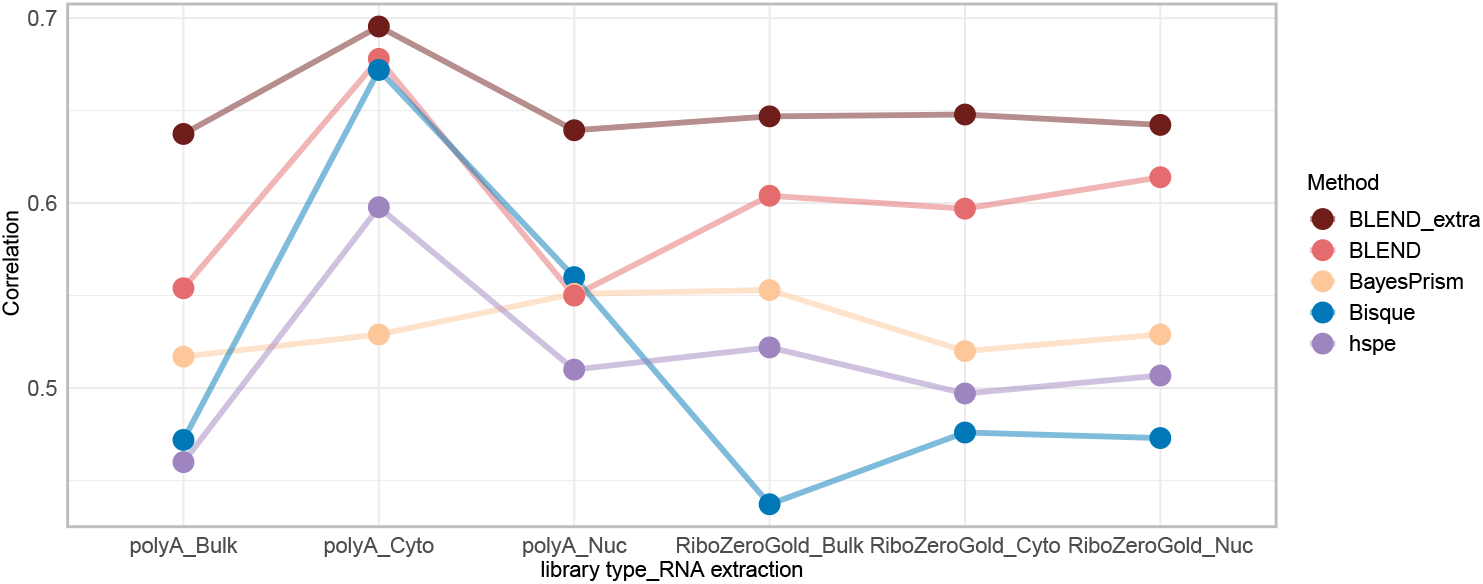
Benchmarking cellular deconvolution methods with real human DLPFC data. Pearson’s correlation coefficients are calculated across cell types and bulk samples under different library types and RNA extraction combinations [17]. MuSiC is not presented because of negative correlations in most settings. BLEND extra is the setting where BLEND additionally used the Sutton reference list.

Furthermore, BLEND’s performance can be improved with richer reference information (Fig. 4). Sutton et al. [21] collected nine brain datasets and provided a list of corresponding CTS reference matrices [22–30] with 6,388 genes. These datasets were from varied sources and sequenced by different technologies. Reference names and original data sources are in Supplementary Notes. Without any data evaluation and processing, we added these references to the reference list and ran BLEND again. These extra references further improved BLEND’s performance across all conditions. BLEND shows high robustness when provided with varied references. The practical implication is that users can simply provide BLEND with all available references without worrying if data with unsatisfying quality will bias the estimation results.

### 2.5 BLEND detects cell type abundance changes in Alzheimer’s disease progression

To demonstrate BLEND’s performance in downstream analyses, we applied BLEND to detect cellular abundance changes in Alzheimer’s disease. The Mount Sinai Brain Bank (MSBB) study [31] provided bulk RNA-seq data of 894 samples collected from postmortem human brain, 850 samples of which have corresponding Braak AD-staging score (bbscore) for progression of neurofibrillary neuropathology. Brodmann area information was also provided. To illustrate the adaptability of BLEND’s automated reference selection, we directly used the Sutton reference list [21].

We visualized cellular fraction changes with boxplots for three major AD-associated cell types: excitatory neurons, inhibitory neurons, and microglia (Figs. 5a-c). We also reported Pearson’s correlation coefficients, identifying a significant decrease in excitatory and inhibitory neurons and an increase in microglia.

**Fig. 5.**
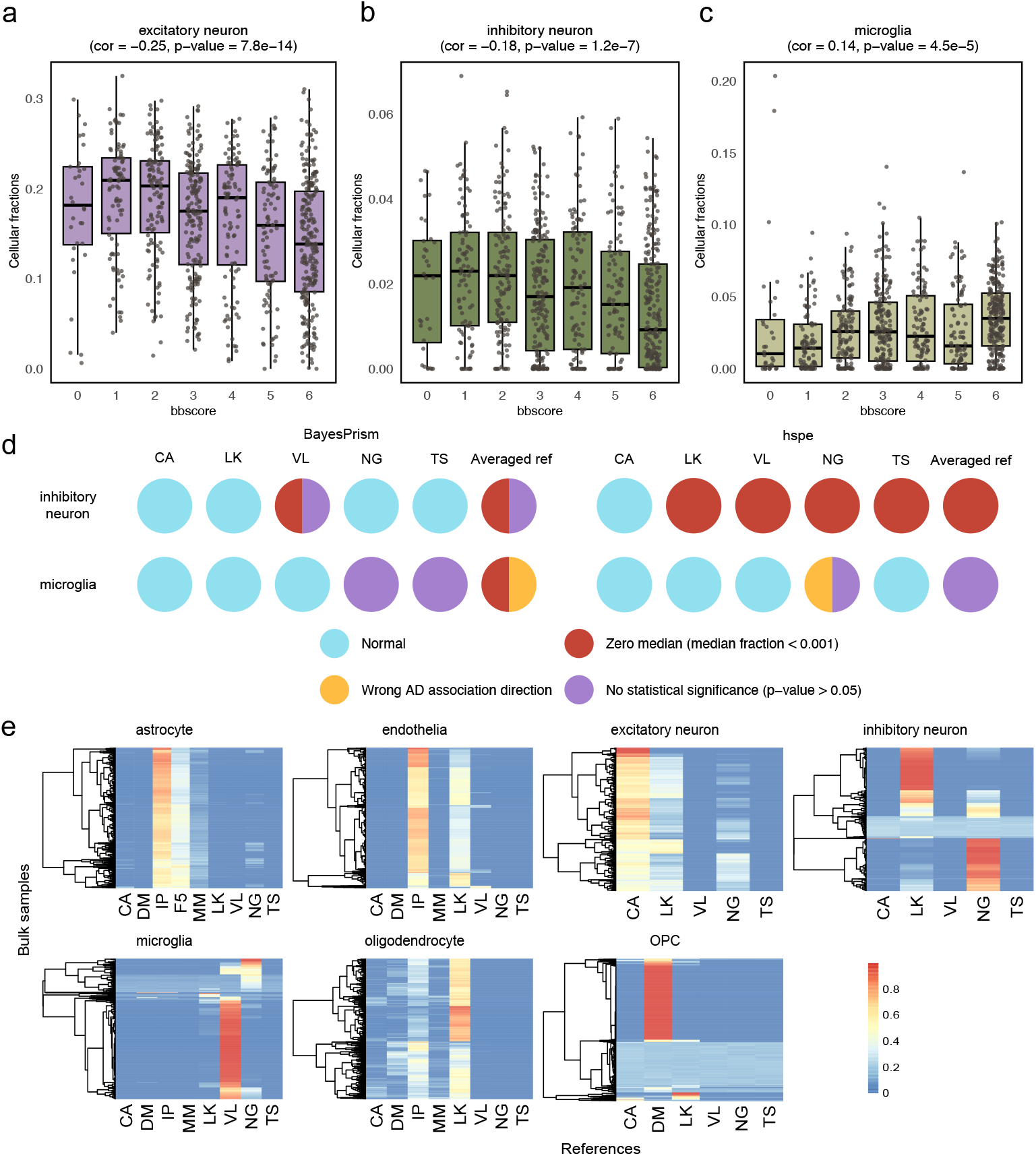
Analyzing MSBB postmortem human brain data. **a-c**, Differential cellular fractions with Alzheimer’s disease progression of three cell types. The x-axes represent Braak AD-staging score (bbscore), and the y-axes represent cellular fractions. Correlation test results between bbscore and cellular fractions are shown in brackets below titles. **d**, Issues of BayesPrism and hspe when being used to repeat differential analyses using different single references and the averaged reference. Colors of pie charts represent issues the corresponding settings have. **e**, Heatmap of reference mixing proportions. Rows represent bulk samples, and columns represent references.

Although these associations can be detected by other methods by using certain references, users may suffer from selective reporting. BayesPrism and hspe allow single reference matrix input. We applied BayesPrism and hspe to deconvolve the MSBB data using single references in the Sutton reference list that have all cell types and the averaged reference. Then, we repeated AD association analyses. Unfortunately, most references suffer from issues including (1) zero median fraction estimates (*<* 0.001), (2) wrong AD association direction, and (3) no statistical significance (p-value *>* 0.05) (Fig. 5d). BLEND helps avoid these issues by performing automated reference selection.

The estimated reference mixing proportions can help us understand the importance of reference selection (Fig. 5d-j). While reference mixing proportions of bulk samples were estimated independently, heatmaps all presented non-random patterns. (1) First, reference quality matters the most. Astrocytes and endothelial cells both preferred the IP reference [28], where authors acutely purified human astrocytes along with other major cell types, serving as a high-quality reference. Excitatory neurons selected CA reference [22], in which only excitatory neurons’ labels were validated using RNAscope mFISH technology and thus have promising quality. OPCs preferred the DM reference [23], derived from surgically resected brain tissues that generally have higher quality than frozen tissues. (2) When two references have similar qualities, some biological factors start to show. Interestingly, for inhibitory neurons, we found a significant association between the Brodmann area of bulk samples and their reference preference (Chi-squared test, p-value*<* 2.2 × 10^−16^). The NG reference [25] measured human brain samples from Brodmann area 9, and 45.4% of bulk samples choosing NG were from the adjacent Brodmann area 10. The LK reference [27] includes information from Brodmann areas responsible for human executive functions, with 44.0% of bulk samples choosing LK from area 44, located in this functional region.

## 3 Discussion

In summary, we presented BLEND, a novel hierarchical Bayesian model for cellular deconvolution that estimates cellular fractions accurately and robustly, without the unrealistic assumptions of no discrepancy reference and uniform CTS gene expression across bulk samples in many existing deconvolution methods. By customizing references for bulk samples through exploring convex hulls of all the available references, BLEND elegantly accommodates cross-sample, cross-data, and cross-technology heterogeneity to improve cellular fraction estimation. We validated BLEND’s performance through comprehensive benchmarking using both simulated and real multi-assay data from postmortem human brain, revealing its superior accuracy and robustness in estimating cellular fractions than state-of-the-art methods. By deconvolving large-scale human brain data, we detected cellular abundance changes with AD progression, showcasing its potential in published data integration and real-life application.

Notably, BLEND demonstrated its capability to find the most suitable references for bulk samples automatically, eliminating the need for reference quality evaluation. Moreover, BLEND needs no data transformation/normalization or cell type marker gene selection before deconvolution. All of these facts illustrate that BLEND is an accurate, robust, and easy-to-use deconvolution tool.

Nevertheless, there are some drawbacks of BLEND. First, the computational time of our Gibbs sampler and EM-MAP algorithm can be relatively long, especially when all genes are used and hundreds of references are provided to BLEND. However, the EM-MAP algorithm has been implemented using Rcpp and allows parallelization, which can accelerate computation enormously. Second, gene-gene correlation is ignored in BLEND for model simplicity. Thus, even though we can estimate the posterior distribution, statistical inference for cellular fractions is not appropriate. This issue may not be major since most downstream analysis tools do not take the uncertainty in point estimates into account. Providing accurate point estimates is more useful than accounting for the uncertainty of biased estimates.

BLEND serves as an accurate and robust tool for cell type-specific analyses. In the future, it will be interesting to design a unified statistical tool that can perform genegene correlation-aware deconvolution utilizing multiple references and incorporate cellular fraction estimate uncertainties in differential analysis.

## 4 Methods

### 4.1 Notation used throughout the manuscript

For any positive integer *n*, we let [*n*] = {1, …, *n*}. For ***p*** ∈ [0, 1]^*n*^ satisfying 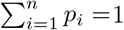, we call the random variable *x* ∈ [*n*] categorically distributed as *x* ∼ Cat([*n*], ***p***) if ℙ(*x* = *i*) = *p*_*i*_ for all *i* ∈ [*n*].

### 4.2 Overview of BLEND’s probability model

BLEND models the deconvolution of bulk count RNA-seq data using multiple CTS gene expression references. Let *X*_*n,g*_ ∈ ℕ_0_ be the bulk RNA-seq count for gene *g* ∈ [*G*] in subject *n* ∈ [*N* ] and 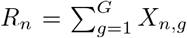 be the read depth for subject *n*. Let ***µ***_*n*_ ∈ [0, 1]^*T*^ be the *n*-th subject’s vector of cell type fractions across *T* cell types, which satisfies 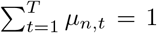. We assume there are *M*_*t*_ available references for cell type *t* and let **Φ**_*t,m*_ ∈ [0, 1]^*G*^ be the *m*-th reference for cell type *t*. As this is a distribution over genes, 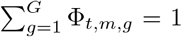. Our goal is to estimate cell type proportions ***µ***_*n*_ given the observed data *X*_*n,g g*∈[*G*]_ and references **Φ**_*t,m t*∈[*T* ];*m*∈[*M t*_ ].

BLEND uses a generative “bag-of-words” model [10] to generate each read’s gene identity. Briefly, we first draw the read’s cell type of origin. The probability the read is assigned to gene *g* is then determined by the originating cell type’s reference. Unlike traditional LDA, which assumes references are the same across subjects [10], BLEND allows references to be subject-specific by assuming that the reference for cell type *t* in subject *n* lies in the convex hull of {**Φ**_*t,m*_}_*m*∈[*Mt*_]. We formalize these ideas in the below generative probability model, which generates the observed data{ *X*_*n,g*_ }_*g*∈[*G*]_ for each subject *n*. The graphical plate representation is in Supplementary Notes. We implicitly condition on read depth *R*_*n*_ and observed references **Φ**_*t,m*_, although we leave them out of conditioning arguments for presentation simplicity.

i. Generate a length-*T* vector of cell type proportions ***µ***_*n*_ | ***α*** ∼ Dirichlet(***α***).
ii. For each cell type *t* ∈ [*T* ], generate a length-*M*_*t*_ vector of reference mixing proportions ***ψ***_*n,t*_ | ***β***_*t*_ ∼ Dirichlet(***β***_*t*_).
iii. For read *r* ∈ [*R*_*n*_], draw its cell type source *Y*_*n,r*_ | ***µ***_*n*_ ∼ Cat([*T* ], ***µ***_*n*_).
iv. Draw the *r*-th read’s gene identity,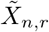, as 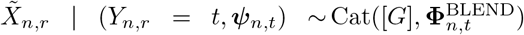, where 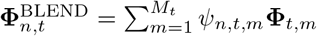.
v. For each gene *g* ∈ [*G*], set 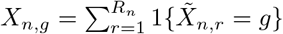.

Step (iv) lets each subject have their own reference 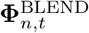 for each cell type *t*, which is assumed to be a convex combination of the observed references {**Φ**_*t,m*_ } _*m*∈[*M t*_ ]. This is quite general, as it allows a subject’s reference to be the average of observed references (*ψ*_*n,t,m*_ = 1*/M*_*t*_ for all *m*) or be determined by only a subset of references (*ψ*_*n,t,m*_ is large for some references *m* and small for others). The latter will occur when references are sampled from multiple populations, where *ψ*_*n,t,m*_ will be large if reference *m* and the CTS expression for subject *n* are sampled from the same population. This ensures our model captures the variation in CTS expression across subjects.

### 4.3 Parameter estimation strategies of BLEND

Here we derive a Gibbs sampler to sample from the posterior ℙ(***µ***_*n*_, {***ψ***_*n,t*_}_*t*∈[*T* ]_ | {*X*_*n,g*_}_*g*∈[*G*]_, ***α***, {***β***_*t*_}_*t*∈[*T* ]_), where our estimator for cell type proportions is then the posterior expectation of ***µ***_*n*_. To ensure Gibbs updates are tractable, we introduce a new latent variable by noting that drawing the *r*-th read’s gene identity in step (iv) of the above generative probability model is equivalent to the following two-step procedure:

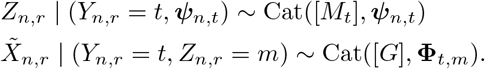

The new latent variable *Z*_*n,r*_ is interpretable as the *r*-th read’s reference source. To derive the Gibbs sampler, let

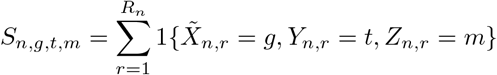

be the number of gene *g* ’s reads that are drawn from cell type *t* using reference *m ∈* [*M*_*t*_], and define ***S***_*n,g*_ = {*S*_*n,g,t,m*_ } _*t*∈[*T* ];*m*∈[*M t*_ ]. In Gibbs sampling, we sample from marginal posteriors

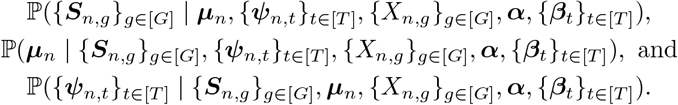

The Gibbs sampler converges quickly due to the large library sizes in bulk RNA-seq data. We let the entries of ***α*** and ***β***_*t*_ be 0.01 in practice, which ensures the priors for ***µ***_*n*_ and ***ψ***_*n,t*_ are non-informative.

While our Gibbs sampler is relatively fast and easy to implement, we would ideally optimize computation to be able to estimate cell-type proportions in modern datasets that include thousands of subjects. We therefore present an algorithm to maximize the posterior distribution, that is, to derive the maximum *a posteriori* (MAP) estimate for ***µ***_*n*_ and {***ψ***_*n,t*_}_*t*∈[*T* ]_. We use an expectation-maximization (EM) algorithm to maximize the log-posterior after introducing the latent variables ***S***_*n,g*_ = *S*_*n,g,t,m t*∈[*T* ];*m*∈[*M t*_ ]. Define the latent variable-augmented log-posterior to be

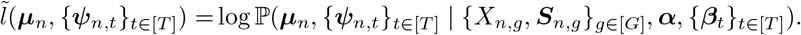

The E-step of the EM algorithm is then:

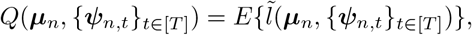

where expectation is taken with respect to the distribution 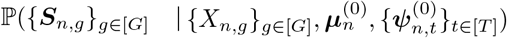 where 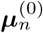 and 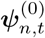 are the algorithm’s current values of ***µ***_*n*_ and ***ψ***_*n,t*_. The M-step is

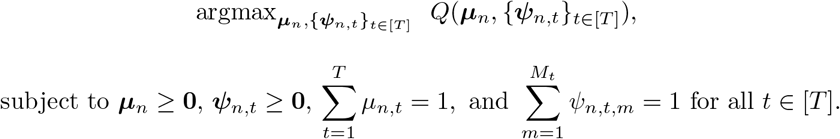

subject to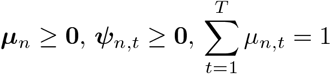, and 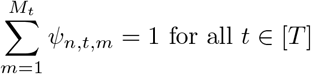.

Aside from being able to provide the same parameter estimates, EM-MAP simplifies estimation by replacing the time-consuming sampling process and the need for additional samples after burn-in with purely algebraic operations and direct parameter estimation upon convergence. We implement the Gibbs sampler in R and the EM-MAP algorithm in Rcpp. In the MSBB application, it takes the Gibbs sampler 36 minutes to deconvolve one bulk sample (6,388 genes) using nine references. Apart from providing consistent estimates (Supplementary Notes), the EM-MAP algorithm significantly reduces computation time to 1.5 minutes.

### 4.4 Cell size adjustment

After the parameter estimation, we discuss the interpretation and use of the cellular fraction parameter ***µ***_*n*_. In BLEND, we model RNA transcripts directly. Thus, ***µ***_*n*_ should be interpreted as proportions of transcript attributed to cell types in the tissue (RNA fractions). When cell fractions are of interest, we need to consider the varying abundance of transcripts of different cell types, which is quantified by a cell size vector 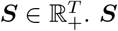 can be given or estimated by the average library sizes of cell types. Then, we estimate cell fractions by

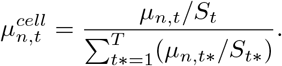

In benchmarking studies with reliable ground truth of cell fractions, such as simulation, we perform cell size adjustment first before comparison.

## Acknowledgements

The results published here are in whole or in part based on data obtained from the AD Knowledge Portal (https://adknowledgeportal.org). Study data were provided by the Rush Alzheimer’s Disease Center, Rush University Medical Center, Chicago. Data collection was supported through funding by NIA grants P30AG10161 (ROS), R01AG15819 (ROSMAP; genomics and RNAseq), R01AG17917 (MAP), R01AG30146, R01AG36042 (5hC methylation, ATACseq), RC2AG036547 (H3K9Ac), R01AG36836 (RNAseq), R01AG48015 (monocyte RNAseq) RF1AG57473 (single nucleus RNAseq), U01AG32984 (genomic and whole exome sequencing), U01AG46152 (ROSMAP AMP-AD, targeted proteomics), U01AG46161(TMT proteomics), U01AG61356 (whole genome sequencing, targeted proteomics, ROSMAP AMP-AD), the Illinois Department of Public Health (ROSMAP), and the Translational Genomics Research Institute (genomic). Additional phenotypic data can be requested at www.radc.rush.edu. The MSBB data were generated from postmortem brain tissue collected through the Mount Sinai VA Medical Center Brain Bank and were provided by Dr. Eric Schadt from Mount Sinai School of Medicine.

## Declarations

## Data availability

This work mainly used five public datasets, including Mathys data (https://www.synapse.org/#!Synapse:syn52293417), Fujita data (https://www.synapse.org/#!Synapse:syn31512863), LIBD data (https://github.com/LieberInstitute/Human_DLPFC_Deconvolution), MSBB bulk RNA-seq data (https://www.synapse.org/Synapse:syn22025006) and Sutton reference list (https://github.com/Voineagulab/BrainCellularComposition).

## Code availability

Package is available at https://github.com/Penghuihuang2000/BLEND.

## Author contribution

Under the supervision of J.W. and C.M., P.H. conceived the idea, derived algorithms, and conducted experiments. P.H., J.W., and C.M. designed the experiments and wrote the manuscript. P.H. and M.C. implemented the software. J.W. provided funding support.

