## Supplementary Notes for "BLEND: Probabilistic Cellular Deconvolution with Automated Reference Selection"

**Huang et al.**

1.5 The Gibbs sampler and the EM-MAP algorithm provide consistent estimates 7

**2 BLEND provides accurate cellular fraction estimates and reference selection** **7**

**3 Sutton reference list** **9**

### 1 Technical details of BLEND

#### 1.1 Limitations of existing methods

Cellular deconvolution has been a popular topic and many methods have been developed in recent years, where bulk data is modeled as the average gene expression of different cell types weighted by their corresponding cellular fractions. Other than statistical models, the performance of deconvolution methods is influenced by several factors:

1. Quality of reference data. In reference-based cellular deconvoluton, methods assume that the underlying CTS gene expression of bulk data is similar to the reference data. As shown in many benchmarking paper, the reference data is the major factor of estimation accuracy.

2. Cell type marker genes. Reference-based methods and marker-based methods are dependent on cell type marker genes for cell type identifiability especially when there are similar cell types to be estimated, e.g., inhibitory neurons and excitatory neurons. Unsupervised methods are also need cell type marker genes to annotate cell types.

3. Data normalization methods, e.g., CPM (counts per million), TPM (transcripts per million), RPKM (reads per kilobase of transcript per million reads mapped), etc.

4. Data transformation methods, e.g., log-fold change or linear.

Recently, there are many open scRNA-seq(sn) data sources. When performing deconvolution, as most methods can only incorporate one reference matrix, the selection or integration of reference datasets still remain unclear. Moreover, methods fail to account for the heterogeneity of CTS gene expression across bulk samples', and the discrepancy between bulk samples and the reference data. How to decide cell type marker genes and the data normalization/transformation methods makes it even harder to use deconvolution methods.

#### 1.2 Model set-up

Based on the aforementioned limitations, it is pressing to develop an easy-to-use new deconvolution method. BLEND is a Bayesian deconvolution method that estimates cellular fractions accurately and robustly by leveraging multiple references, requiring no marker gene selection and data normalization/transformation. BLEND personalizes references for each bulk sample and employs a bag-of-words representation for bulk samples for deconvolution.

For any positive integer $n$, we let $[n] = \{1, \ldots, n\}$. For $\boldsymbol{p} \in [0,1]^n$ satisfying $\sum_{i=1}^{n} p_i = 1$, we say the random variable $x \in [n]$ is categorically distributed as $x \sim \text{Cat}([n], \boldsymbol{p})$ if $\mathbb{P}(x = i) = p_i$ for all $i \in [n]$. Mult is short for multinomial distribution.

Notations that will be used throughout the derivation are as follows:

1. Individual/Subject: $n \in [N]$;

2. Gene: $g \in [G]$;

3. Reference for cell type $t$: $m \in [M_t]$;

4. Cell type: $t \in [T]$;

5. Library size of subject $n$: $R_n$;

6. Read count of individual $n$: $r \in [R_n]$;

7. Bulk counts matrix: $\boldsymbol{X} \in \mathbb{N}_0^{N \times G}$;

8. Cellular fraction vector of individual $n$: $\boldsymbol{\mu}_n \in [0,1]^T$;

9. Reference mixing proportion vector for individual $n$'s cell type $t$: $\boldsymbol{\psi}_{n,t} \in [0,1]^{M_t}$;

10. The $m$th CTS reference for cell type $t$: $\boldsymbol{\Phi}_{t,m} \in [0,1]^G$.

The generative process that generates the observed data $\{X_{n,g}\}_{g \in [G]}$ for each subject $n$ is visually presented in Figure S1a and formalized as follows.

(i) Generate a length-$T$ vector of cell type proportions $\boldsymbol{\mu}_n \mid \boldsymbol{\alpha} \sim \text{Dirichlet}(\boldsymbol{\alpha})$.

(ii) For each cell type $t \in [T]$, generate a length-$M_t$ vector of reference mixing proportions $\boldsymbol{\psi}_{n,t} \mid \boldsymbol{\beta}_t \sim \text{Dirichlet}(\boldsymbol{\beta}_t)$.

(iii) For read $r \in [R_n]$, draw its cell type source $Y_{n,r} \mid \boldsymbol{\mu}_n \sim \text{Cat}([T], \boldsymbol{\mu}_n)$.

(iv) Draw the $r$-th read's gene identity, $\tilde{X}_{n,r}$, as $\tilde{X}_{n,r} \mid (Y_{n,r} = t, \boldsymbol{\psi}_{n,t}) \sim \text{Cat}([G], \boldsymbol{\Phi}_{n,t}^{\text{BLEND}})$, where $\boldsymbol{\Phi}_{n,t}^{\text{BLEND}} = \sum_{m=1}^{M_t} \psi_{n,t,m} \boldsymbol{\Phi}_{t,m}$.

(v) For each gene $g \in [G]$, set $X_{n,g} = \sum_{r=1}^{R_n} 1\{\tilde{X}_{n,r} = g\}$.

**Notes**

- The mixed reference $\boldsymbol{\Phi}_{n,t}^{BLEND}$ is subject-specific and cell type-specific. This allows each subject to have its own CTS personalized reference. Moreover, this design considers the situation where different cell types in the same dataset have varied quality.

- $\boldsymbol{\Phi}_{n,t}^{\text{BLEND}} = \sum_{m=1}^{M_t} \psi_{n,t,m} \boldsymbol{\Phi}_{t,m}$ can be any vector in the convex hull of available references $\{\ \boldsymbol{\Phi}_{t,m}\}_{m \in [M_t]}$, which is rather flexible. For example, it allows a subject's reference to be the average of observed references ($\psi_{n,t,m} = 1/M_t$ for all $m$) or be determined by only a subset of references ($\psi_{n,t,m}$ is large for some references $m$ and small for others). The latter will occur when references are sampled from multiple populations, where $\psi_{n,t,m}$ will be large if reference $m$ and the CTS expression for subject $n$ are sampled from the same population.

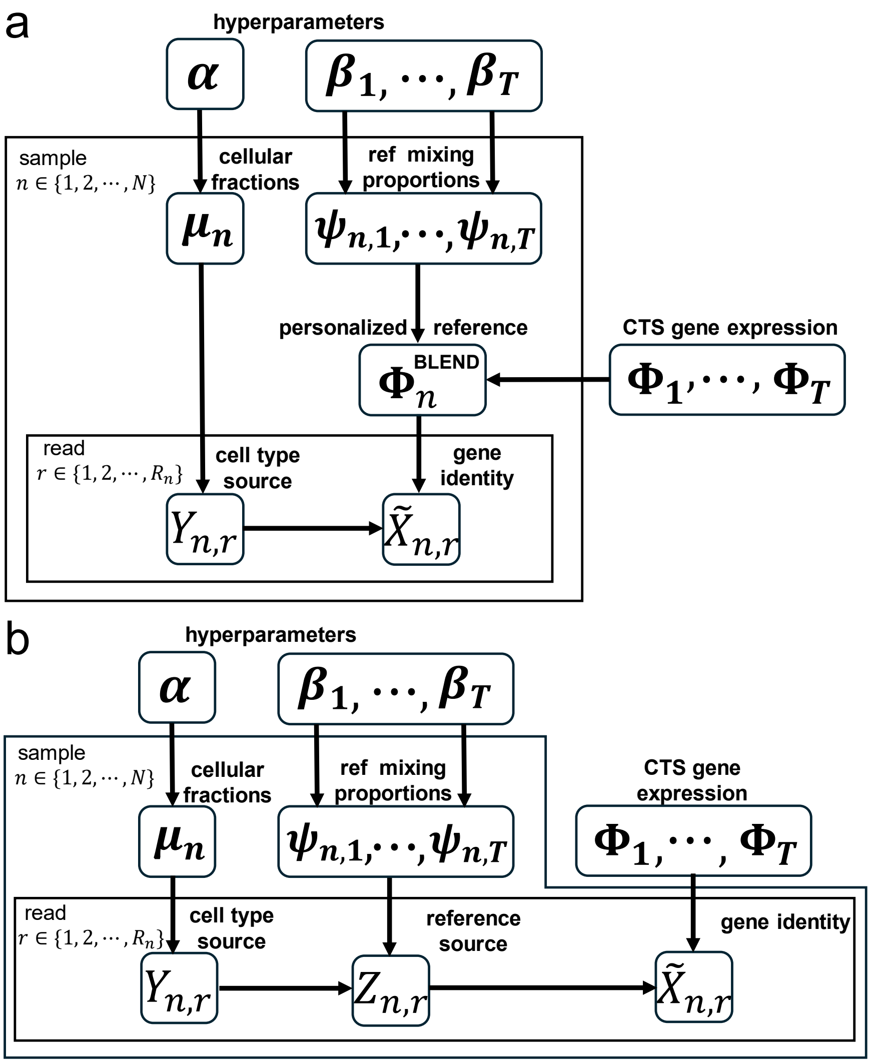

Figure S1: Plate representations of BLEND. (a) The original plate representation of BLEND. (b) The equivalent plate representation of BLEND after introducing the latent variable $Z_{n,r}$.

#### 1.3 Derivation of the Gibbs sampler

To derive the Gibbs sampler, we introduce a latent variable by noting drawing the $r$-th read's gene identity in step (iv) of the above generative probability model is equivalent to the following two-step procedure:

$$Z_{n,r} \mid (Y_{n,r} = t, \boldsymbol{\psi}_{n,t}) \sim \text{Cat}([M_t], \boldsymbol{\psi}_{n,t})$$
$$\tilde{X}_{n,r} \mid (Y_{n,r} = t, Z_{n,r} = m) \sim \text{Cat}([G], \boldsymbol{\Phi}_{t,m}).$$

The new latent variable $Z_{n,r}$ is interpretable as the $r$-th read's reference source and present the equivalent plate representation of BLEND with $Z_{n,r}$ in Figure S1b. The hierarchical model is then:

(i) $\boldsymbol{\mu}_n \mid \boldsymbol{\alpha} \sim \text{Dirichlet}(\boldsymbol{\alpha})$.

(ii) $\boldsymbol{\psi}_{n,t} \mid \boldsymbol{\beta}_t \sim \text{Dirichlet}(\boldsymbol{\beta}_t)$.

(iii) $Y_{n,r} \mid \boldsymbol{\mu}_n \sim \text{Cat}([T], \boldsymbol{\mu}_n)$.

(iv) $Z_{n,r} \mid (Y_{n,r} = t, \boldsymbol{\psi}_{n,t}) \sim \text{Cat}([M_t], \boldsymbol{\psi}_{n,t})$.

(v) $\tilde{X}_{n,r} \mid (Y_{n,r} = t, Z_{n,r} = m) \sim \text{Cat}([G], \boldsymbol{\Phi}_{t,m})$.

(vi) For each gene $g \in [G]$, set $X_{n,g} = \sum_{r=1}^{R_n} 1\{\tilde{X}_{n,r} = g\}$.

To derive the Gibbs sampler, let

$$S_{n,g,t,m} = \sum_{r=1}^{R_n} 1\{\tilde{X}_{n,r} = g, Y_{n,r} = t, Z_{n,r} = m\}$$

be the number of gene $g$'s reads that are drawn from cell type $t$ using reference $m \in [M_t]$, and define $\boldsymbol{S}_{n,g} = \{S_{n,g,t,m}\}_{t\in[T];m\in[M_t]}$. We let $\boldsymbol{\mu}$, $\boldsymbol{\psi}$, $\tilde{\boldsymbol{X}}$, $\boldsymbol{Y}$ and $\boldsymbol{Z}$ denote $\{\boldsymbol{\mu}_n\}_{n\in[N]}$, $\{\boldsymbol{\psi}_{n,t}\}_{n\in[N],t\in[T]}$ ,$\{\tilde{X}_{n,r}\}_{n\in[N],r\in[R_n]}$, $\{Y_{n,r}\}_{n\in[N],r\in[R_n]}$ and $\{Z_{n,r}\}_{n\in[N],r\in[R_n]}$, respectively.

Here is the detailed derivation.

$$\begin{aligned}
&\mathbb{P}(\boldsymbol{\mu}_n \mid \boldsymbol{\mu}_{-n}, \boldsymbol{\psi}, \tilde{\boldsymbol{X}}, \boldsymbol{Y}, \boldsymbol{Z}, \boldsymbol{X}) \\
&\propto \mathbb{P}(\boldsymbol{X}_n \mid \boldsymbol{\mu}_n, \boldsymbol{\mu}_{-n}, \boldsymbol{\psi}, \tilde{\boldsymbol{X}}_n, \boldsymbol{Y}_n, \boldsymbol{Z}_n)\mathbb{P}(\boldsymbol{\mu}_n \mid \boldsymbol{\mu}_{-n}, \boldsymbol{\psi}, \tilde{\boldsymbol{X}}_n, \boldsymbol{Y}_n, \boldsymbol{Z}_n) \\
&\propto \mathbb{P}(\boldsymbol{\mu}_n \mid \boldsymbol{\mu}_{-n}, \boldsymbol{\psi}, \tilde{\boldsymbol{X}}_n, \boldsymbol{Y}_n, \boldsymbol{Z}_n) \\
&\propto \mathbb{P}(\boldsymbol{Y}_n \mid \boldsymbol{\mu}_n)\mathbb{P}(\boldsymbol{\mu}_n) \\
&\propto \prod_{t=1}^{T} \mu_{n,t}^{\alpha_t - 1} \prod_{r=1}^{R_n} \mu_{n,t}^{1\{Y_{n,r}=t\}} \\
&= \prod_{t=1}^{T} \mu_{n,t}^{\alpha_t + \sum_{m=1}^{M_t} \sum_{g=1}^{G} S_{n,g,t,m} - 1}
\end{aligned}$$

$$\begin{aligned}
&\mathbb{P}(\boldsymbol{\psi}_{n,t} \mid \boldsymbol{\psi}_{-(n,t)}, \boldsymbol{\mu}, \tilde{\boldsymbol{X}}, \boldsymbol{Y}, \boldsymbol{Z}, \boldsymbol{X}) \\
&\propto \mathbb{P}(\boldsymbol{X}_n \mid \boldsymbol{\psi}_{n,t}, \boldsymbol{\psi}_{-(n,t)}, \boldsymbol{\mu}, \tilde{\boldsymbol{X}}, \boldsymbol{Y}, \boldsymbol{Z})\mathbb{P}(\boldsymbol{\psi}_{n,t} \mid \boldsymbol{\psi}_{-(n,t)}, \boldsymbol{\mu}, \tilde{\boldsymbol{X}}, \boldsymbol{Y}, \boldsymbol{Z}) \\
&\propto \mathbb{P}(\boldsymbol{Z}_n \mid \boldsymbol{\psi}_{n,t}, \boldsymbol{Y}_n)\mathbb{P}(\boldsymbol{\psi}_{n,t}) \\
&\propto \prod_{m=1}^{M_t} \psi_{n,t,m}^{\beta_{t,m} - 1} \prod_{r=1}^{R_n} \psi_{n,t,m}^{1\{\ Y_{n,r}=t, Z_{n,r}=m\}} \\
&= \prod_{m=1}^{M_t} \psi_{n,t,m}^{\beta_{t,m} + \sum_{g=1}^{G} S_{n,g,t,m} - 1}
\end{aligned}$$

$$\begin{aligned}
&\mathbb{P}(\boldsymbol{S}_{n,g} \mid \boldsymbol{S}_{-(n,g)}, \boldsymbol{\mu}, \boldsymbol{\psi}, \boldsymbol{X}) \\
&= \mathbb{P}(\tilde{\boldsymbol{X}}_n, \boldsymbol{Y}_n, \boldsymbol{Z}_n \mid \tilde{\boldsymbol{X}}_{-n}, \boldsymbol{Y}_{-n}, \boldsymbol{Z}_{-n}, \boldsymbol{\mu}_n, \boldsymbol{\psi}_n, \boldsymbol{X})
\end{aligned}$$

$$
\begin{aligned}
&= \mathbb{P}(\tilde{\boldsymbol{X}}_n \mid \boldsymbol{Y}_n, \boldsymbol{Z}_n, \boldsymbol{\mu}_n, \boldsymbol{\psi}_n, \boldsymbol{X})\mathbb{P}(\boldsymbol{Z}_n \mid \boldsymbol{Y}_n, \boldsymbol{\mu}_n, \boldsymbol{\psi}_n, \boldsymbol{X})\mathbb{P}(\boldsymbol{Y}_n \mid \boldsymbol{\mu}_n, \boldsymbol{\psi}_n, \boldsymbol{X}) \\
&\propto (\prod_{t=1}^{T}\prod_{m=1}^{M_t}\prod_{g=1}^{G} \Phi_{t,m,g}^{\sum_{r=1}^{R_n} 1\{\tilde{X}_{n,r}=g, Y_{n,r}=t, Z_{n,r}=m\}})(\prod_{t=1}^{T}\prod_{m=1}^{M_t} \psi_{n,t,m}^{\sum_{r=1}^{R_n} 1\{Y_{n,r}=t, Z_{n,r}=m\}}) \\
&\quad (\prod_{t=1}^{T} \mu_{n,t}^{\sum_{r=1}^{R_n} 1\{Y_{n,r}=t\}}) \\
&= \prod_{t=1}^{T}\prod_{m=1}^{M_t}\prod_{g=1}^{G} (\mu_{n,t}\psi_{n,t,m}\Phi_{t,m,g})^{S_{n,g,t,m}}
\end{aligned}
$$

Then, it yields the Gibbs sampling steps:

$$
\begin{aligned}
&\boldsymbol{S}_{n,g} \mid \cdot \sim \text{Mult}(X_{n,g}, \boldsymbol{P}_{n,g}), \quad P_{n,g,t,m} \propto \mu_{n,t}\psi_{n,t,m}\Phi_{t,m,g}, \quad g \in [G]; t \in [T]; m \in [M_t] \\
&\boldsymbol{\psi}_{n,t} \mid \cdot \sim \text{Dirichlet}(\beta_{t,1} + \textstyle\sum_{g=1}^{G} S_{n,g,t,1}, \ldots, \beta_{t,M_t} + \sum_{g=1}^{G} S_{n,g,t,M_t}), \quad t \in [T] \\
&\boldsymbol{\mu}_n \mid \cdot \sim \text{Dirichlet}(\alpha_1 + \textstyle\sum_{g=1}^{G}\sum_{m=1}^{M_1} S_{n,g,t,m}, \ldots, \alpha_T + \sum_{g=1}^{G}\sum_{m=1}^{M_T} S_{n,g,T,m}).
\end{aligned}
$$

#### 1.4 Derivation of the EM-MAP algorithm

$$
\begin{aligned}
\tilde{l}(\boldsymbol{\mu}_n, \{\boldsymbol{\psi}_{n,t}\}_{t\in[T]}) &= \log \mathbb{P}(\boldsymbol{\mu}_n, \{\boldsymbol{\psi}_{n,t}\}_{t\in[T]} \mid \{X_{n,g}, \boldsymbol{S}_{n,g}\}_{g\in[G]}, \boldsymbol{\alpha}, \{\boldsymbol{\beta}_t\}_{t\in[T]}) \\
&= \log \mathbb{P}(\boldsymbol{\mu}_n, \{\boldsymbol{\psi}_{n,t}\}_{t\in[T]} \mid \{\boldsymbol{S}_{n,g}\}_{g\in[G]}, \boldsymbol{\alpha}, \{\boldsymbol{\beta}_t\}_{t\in[T]}) \\
&= \sum_{g=1}^{G}\sum_{t=1}^{T}\sum_{m=1}^{M_t} \log \mathbb{P}(S_{n,g,t,m} \mid \mu_{t,n}, \psi_{n,t,m}) + \log \mathbb{P}(\boldsymbol{\mu}_n \mid \boldsymbol{\alpha}) + \sum_{t=1}^{T} \log \mathbb{P}(\boldsymbol{\psi}_{n,t} \mid \boldsymbol{\beta}_t)
\end{aligned}
$$

The E-step is

$$
\begin{aligned}
&Q(\boldsymbol{\mu}_n, \{\boldsymbol{\psi}_{n,t}\}_{t\in[T]}) = E\{\tilde{l}(\boldsymbol{\mu}_n, \{\boldsymbol{\psi}_{n,t}\}_{t\in[T]})\} \\
&= \sum_{g=1}^{G}\sum_{t=1}^{T}\sum_{m=1}^{M_t} U_{n,g,t,m}(\log \mu_{n,t}\Phi_{t,m,g}\psi_{n,t,m}) + \sum_{t=1}^{T}(\alpha_t - 1)\log \mu_t + \sum_{t=1}^{T}\sum_{m=1}^{M_t}(\beta_{t,m} - 1)\log \psi_{n,t,m}.
\end{aligned}
$$

The M-step is

$$
\text{argmax}_{\boldsymbol{\mu}_n, \{\boldsymbol{\psi}_{n,t}\}_{t\in[T]}} \; Q(\boldsymbol{\mu}_n, \{\boldsymbol{\psi}_{n,t}\}_{t\in[T]}),
$$

$$
\text{subject to } \boldsymbol{\mu}_n \geq \mathbf{0},\ \boldsymbol{\psi}_{n,t} \geq \mathbf{0},\ \sum_{t=1}^{T} \mu_{n,t} = 1, \text{ and } \sum_{m=1}^{M_t} \psi_{n,t,m} = 1 \text{ for all } t \in [T].
$$

To solve this, we introduce Lagrangian function:

$$
Q^* = Q + \lambda_1(1 - \sum_{t=1}^{T} \mu_t) + \sum_{t=1}^{T} \lambda_{2,t}(1 - \sum_{m=1}^{M_t} \psi_{n,t,m}) - \sum_{t=1}^{T} \eta_{1,t}\mu_{n,t} - \sum_{t=1}^{T}\sum_{m=1}^{M_t} \eta_{2,t,m}\psi_{n,t,m}.
$$

By applying the Karush–Kuhn–Tucker (KKT) conditions, we get the algorithm in main text Algorithm 1.

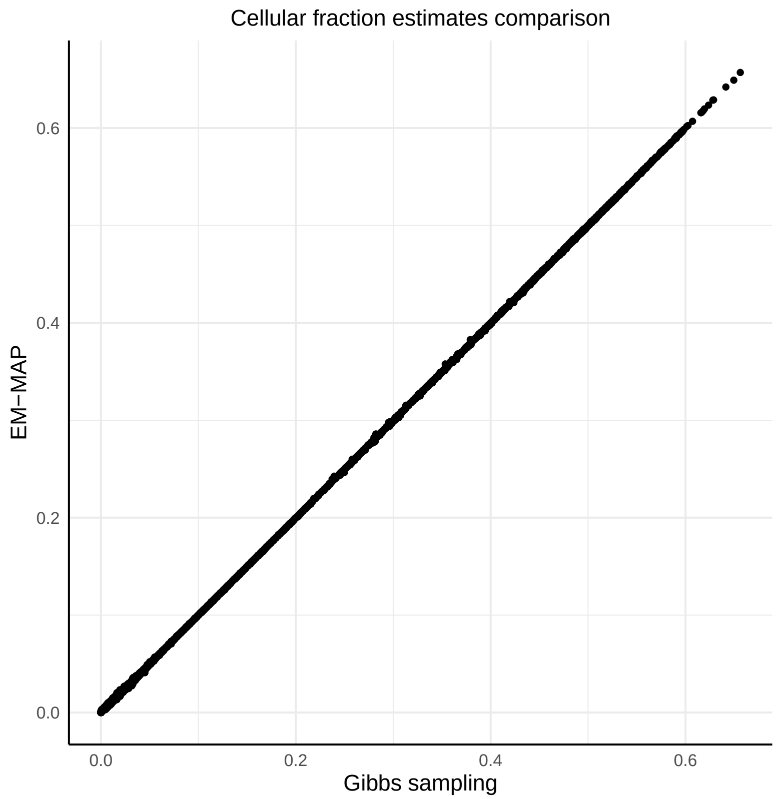

Figure S2: **Comparison between cellular fraction estimates from the Gibbs sampler and the EM-MAP algorithm.** Two algorithms were used to deconvolve MSBB brain data using the Sutton reference list. The x-axis represents estimates from the Gibbs sampler and the y-axis represents estimates from the EM-MAP algorithm.

##### 1.5 The Gibbs sampler and the EM-MAP algorithm provide consistent estimates

Gibbs sampling and EM-MAP are two approaches to estimate parameters of the posterior distribution. Thus, they should provide consistent estimates. To illustrate our algorithms, we ran the Gibbs sampler and EM-MAP algorithm on the MSBB brain data (M. Wang et al., 2018) that were used in the main text with Sutton reference list (Sutton et al., 2022). Cellular fraction estimates from both algorithms were visualized in Figure S2. Moreover, it takes the Gibbs sampler 36 minutes to deconvolve one bulk sample (6,388 genes). Apart from providing consistent estimates, the EM-MAP algorithm significantly reduces computation time to 1.5 minutes.

#### 2 BLEND provides accurate cellular fraction estimates and reference selection

We first evaluated BLEND's ability to select references by simulating bulk data from a real human brain single nucleus RNA-seq (snRNA-seq) dataset. We designed two numerical experiments: 1) "reference selection", in which each simulated bulk sample's corresponding matched reference was provided but needed to be identified; and 2) "reference averaging", in which a bulk samples' CTS gene expression was the average of the provided references, which BLEND needed to learn.

Mathys et al. (Mathys et al., 2023) collected 427 individuals' postmortem dorsolateral prefrontal cortex (DLPFC) tissues from Religious Orders Study and Rush Memory and Aging Project (ROSMAP) (Bennett et al., 2018). They measured gene expression levels across 2.3 million nuclei isolated from these tissues using droplet-based snRNA-seq. Among all donors, 418 of them contained cells of all six major cell types in the brain: astrocyte (astro), microglia (immune), inhibitory neuron (inh), oligodendrocyte progenitor cell (OPC), oligodendrocyte (oligo) and excitatory neuron (ex). For illustration, we randomly picked 20 individuals' snRNA-seq data ($M = 20$) from the Mathys dataset to simulate 100 bulk samples ($N = 100$) with varying cellular fractions. The detailed data generative process to simulate bulk data is described in Methods. Briefly, we first calculated the CTS gene expression of 20 individuals. In the "reference selection" experiment, each simulated bulk sample's CTS gene expression was one of the 20 individuals' CTS expression. Each individual was used to simulate five bulk samples ($K = 5, N = M \times K$). In "reference averaging", all simulated bulk samples' CTS gene expression was the average CTS expression across 20 individuals. Notably, BLEND was unaware of the simulation setting and needed to learn the CTS expression for each bulk sample. Simulated bulk samples had varying library sizes and cellular fractions (Methods).

Specifically, using individual $m$'s snRNA-seq data, we calculated CTS gene expression $\boldsymbol{\Phi}_{t,m}$, satisfying $\sum_{g=1}^{G} \Phi_{t,m,g} = 1, t \in \{1, \cdots, 6\}$, by simply collapsing cells from cell type $t$ and normalizing the total expression of each cell type to have a sum of one. Cellular fractions $\boldsymbol{\mu}_n$ of simulated bulk sample $n$ were sampled from Dirichlet$(1, 2, 3, 4, 5, 6)$ and its library size $R_n$ was sampled from Uniform$(18,000,000,\ 22,000,000)$, which was a common library size for bulk RNA-seq (Conesa et al., 2016). In reference selection simulation, individual snRNA-seq data $m$ was used to simulate 5 bulk samples ($K = 5, N = M \times K$) whose reference mixing proportions $\boldsymbol{\psi}_{n,t} = (0, \cdots, 1, \cdots, 0)'$ (1 on the $m$th entry). In reference exploration simulation, all the 100 bulk samples had $\boldsymbol{\psi}_{n,t} = (1/M)\mathbf{1}'$. Then, following BLEND's data generative process, we generated the simulated bulk data of two simulation settings by $\boldsymbol{X}_n = \text{round}(R_n \sum_{t=1}^{6} \mu_n^k \sum_{m=1}^{M} \psi_{n,t,m} \boldsymbol{\Phi}_{t,m,\cdot})$.

We calculated mean reference mixing proportions $\bar{\hat{\boldsymbol{\psi}}}_n = \sum_{t=1}^{T} \hat{\boldsymbol{\psi}}_{n,t}/T$ across cell types for each bulk sample. We visualized $\bar{\hat{\boldsymbol{\psi}}}_n$ in reference selection simulation by a heatmap, where we sorted bulk samples and references by individuals. We found that the heatmap presented a block diagonal pattern (Figure S3a), and the best reference proportions had a median of 0.96 (Figure S3b). In the reference exploration simulation, the heatmap showed a uniform pattern (Figure S3c), and the median proportion of references was, as expected, $1/20 = 0.05$ (Figure S3d).

Considering the high similarity among the provided references (average Pearson's correlation = 0.92), it was encouraging that BLEND could distinguish them and learn reference patterns accurately. The cellular fraction estimates were well-estimated, with mean absolute error smaller than 0.001.

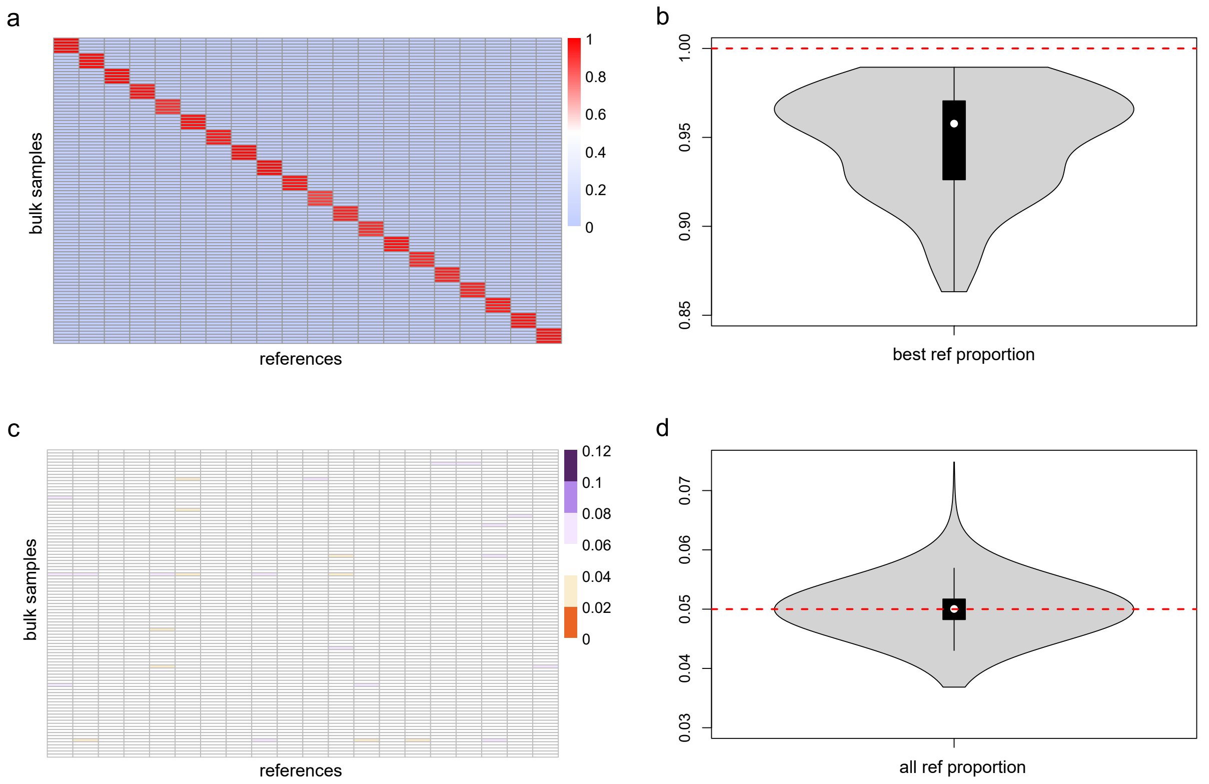

Figure S3: **BLEND enables accurate reference selection.** **a**, Reference selection, in which each bulk sample has its best reference provided in data. Heatmap of the average of reference mixing proportions across cell types. Bulk samples and references are sorted by individuals. **b**, Violin plot of the diagonal elements in the heatmap, which are the proportions of selected best references for bulk samples. **c**, Reference exploration, in which all bulk samples' CTS expression is a uniform combination of all references. Heatmap of mean estimated proportions (averaged across cell types). **d**, Violin plot of mean estimated reference proportions.

### 3 Sutton reference list

| **Name** | **Reference** |
|---|---|
| CA | Hodge, Rebecca D., Trygve E. Bakken, Jeremy A. Miller, Kimberly A. Smith, Eliza R. Barkan, Lucas T. Graybuck, Jennie L. Close et al. "Conserved cell types with divergent features in human versus mouse cortex." *Nature* 573, no. 7772 (2019): 61-68. |
| DM | Darmanis, Spyros, Steven A. Sloan, Ye Zhang, Martin Enge, Christine Caneda, Lawrence M. Shuer, Melanie G. Hayden Gephart, Ben A. Barres, and Stephen R. Quake. "A survey of human brain transcriptome diversity at the single cell level." *Proceedings of the National Academy of Sciences* 112, no. 23 (2015): 7285-7290. |

| | |
|---|---|
| F5 | The FANTOM Consortium and the RIKEN PMI and CLST (DGT). "A promoter-level mammalian expression atlas." *Nature* 507, no. 7493 (2014): 462-470. |
| NG | Nagy, Corina, Malosree Maitra, Arnaud Tanti, Matthew Suderman, Jean-Francois Théroux, Maria Antonietta Davoli, Kelly Perlman et al. "Single-nucleus transcriptomics of the prefrontal cortex in major depressive disorder implicates oligodendrocyte precursor cells and excitatory neurons." *Nature neuroscience* 23, no. 6 (2020): 771-781. |
| VL | Velmeshev, Dmitry, Lucas Schirmer, Diane Jung, Maximilian Haeussler, Yonatan Perez, Simone Mayer, Aparna Bhaduri, Nitasha Goyal, David H. Rowitch, and Arnold R. Kriegstein. "Single-cell genomics identifies cell type–specific molecular changes in autism." *Science* 364, no. 6441 (2019): 685-689. |
| LK | Lake, Blue B., Song Chen, Brandon C. Sos, Jean Fan, Gwendolyn E. Kaeser, Yun C. Yung, Thu E. Duong et al. "Integrative single-cell analysis of transcriptional and epigenetic states in the human adult brain." *Nature biotechnology* 36, no. 1 (2018): 70-80. |
| IP | Zhang, Ye, Steven A. Sloan, Laura E. Clarke, Christine Caneda, Colton A. Plaza, Paul D. Blumenthal, Hannes Vogel et al. "Purification and characterization of progenitor and mature human astrocytes reveals transcriptional and functional differences with mouse." *Neuron* 89, no. 1 (2016): 37-53. |
| MM | Zhang, Ye, Kenian Chen, Steven A. Sloan, Mariko L. Bennett, Anja R. Scholze, Sean O'Keeffe, Hemali P. Phatnani et al. "An RNA-sequencing transcriptome and splicing database of glia, neurons, and vascular cells of the cerebral cortex." *Journal of neuroscience* 34, no. 36 (2014): 11929-11947. |
| TS | Tasic, Bosiljka, Zizhen Yao, Lucas T. Graybuck, Kimberly A. Smith, Thuc Nghi Nguyen, Darren Bertagnolli, Jeff Goldy et al. "Shared and distinct transcriptomic cell types across neocortical areas." *Nature* 563, no. 7729 (2018): 72-78. |
